# Molecular and phylogenetic characterization of the monkeypox outbreak in the South of Spain

**DOI:** 10.1101/2023.09.20.558741

**Authors:** Carlos S. Casimiro-Soriguer, Javier Perez-Florido, Maria Lara, Pedro Camacho-Martinez, Laura Merino-Diaz, Inmaculada Pupo-Ledo, Adolfo de Salazar, Ana Fuentes, Laura Viñuela, Natalia Chueca, Luis Martinez-Martinez, Nicola Lorusso, The Andalusian genomic surveillance network, Jose A Lepe, Joaquín Dopazo, Federico Garcia

## Abstract

Until the May 2022 Monkeypox outbreak, which spread rapidly to many non-endemic countries, the virus was considered a viral zoonosis limited to some African countries. The Andalusian circuit of genomic surveillance was rapidly applied to characterize the Monkeypox outbreak in the South of Spain. Whole genome sequencing was used to obtain the genomic profiles of samples collected across the south of Spain, representative of all the provinces of Andalusia. Phylogenetic analysis was used to study the relationship of the isolates and the available sequences of the 2022 outbreak. Whole genome sequencing of a total of 160 monkeypox viruses from the different provinces that reported cases were obtained. Interestingly, we report the sequences of monkeypox viruses obtained from two patients who died. While one of the isolates bore no noteworthy mutations that explain a potential heightened virulence, in another patient the second consecutive genome sequence, performed after the administration of tecovirimat, uncovered a mutation within the A0A7H0DN30 gene, known to be a prime target for tecovirimat in its Vaccinia counterpart. In general, a low number of mutations were observed in the sequences reported, which were very similar to the reference of the 2022 outbreak (OX044336), as expected from a DNA virus. The samples likely correspond to several introductions of the circulating monkeypox viruses from the last outbreak. The virus sequenced from one of the two patients that died presented a mutation in a gene that bears potential connections to drug resistance. This mutation was absent in the initial sequencing prior to treatment.

## Introduction

Monkeypox (MPXV) is a viral zoonosis endemic in some West and Central African countries and with few cases outside Africa. In May 2022, an unexpectedly large MPXV clade B.1 outbreak affecting a considerable number of non-endemic countries was reported [1, 2]. After the first autochthonous cases were reported in the UK on May 13th and in Spain on May 17th, a rapid spread to more than 14,000 cases were reported in more than 60 countries only in the first two months of the outbreak, that summed up to more than 88,000 cases worldwide as of June 2023 [3]. Although the incidence has drastically reduced in 2023 [3], the observed simultaneous MPXV incidence in different countries due to a rapid cross-border transmission [4] poses a real threat that must be addressed by robust public health surveillance and control measures [5]. Moreover, further research is required to delve deeper into the origins of the recent outbreak, investigating potential factors such as animal reservoirs, human behavior, or viral mutations that might be driving its occurrence. [5]. In this context, genomic monitoring of the MPXV samples sequenced from the epidemiologic surveillance in Andalusia results crucial from the epidemiological point of view, that assigns clearly the Andalusian sequences to the circulating clade. Moreover, whole-genome virus sequencing has also a relevant role in monitoring polymorphisms, as well as in detecting gene losses based on possible intragenic frameshifts or premature stop codons that could appear locally and might be relevant as virulence or enhanced transmissibility determinants.

## Results

### The Andalusian outbreak

The phylogeny of the 2022 MPXV outbreak in the context of the rest of available MPXV sequences, as displayed in the Genomic Surveillance Circuit of Andalusia [6, 7], is depicted in Figure 1. The specific features of the sequences of this outbreak and the apparently fast evolution with respect to previous outbreaks has already been discussed [8]. As expected from a DNA virus, with a relatively low introduction into the general population, the samples isolated in Andalusia have only a few mutations with respect to the rest of MPXV isolates in the outbreak. Andalusian isolates are scattered across the outbreak branch in the phylogeny and are related to sequences from other countries, suggesting different introductions of the virus in Andalusia.

**Figure 1.**
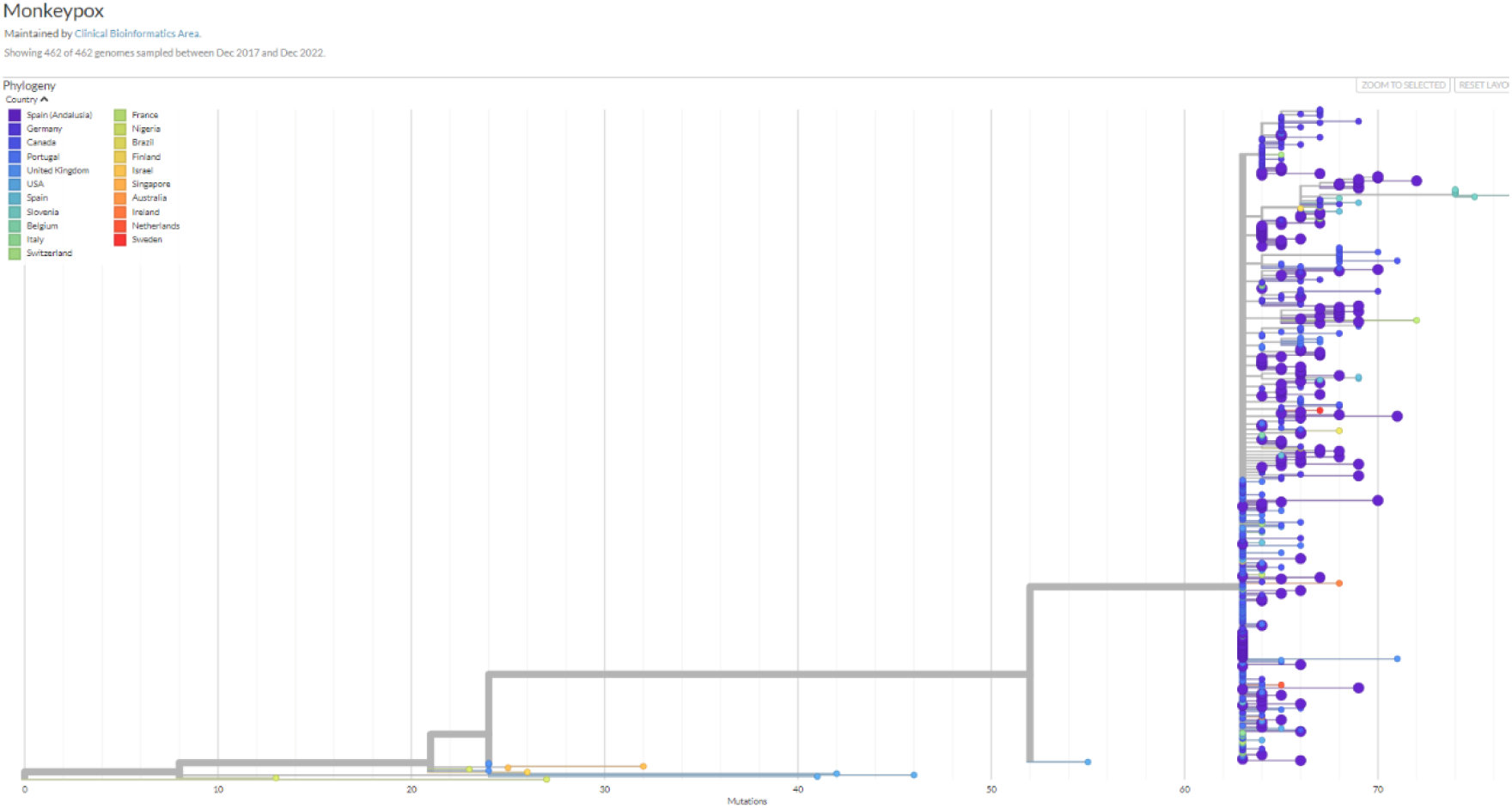
Phylogeny obtained with Nexstrain of the 2022 MPXV outbreak along with the rest of MPXV sequences available.

### Mutational spectrum of the Andalusian outbreak

Figure 2 portrays a detail of the phylogeny of the MPXV isolated in Andalusia (See Supplementary Figure S1 for a more detailed picture). In addition, it is worth analyzing in detail some non-synonymous mutations that appear specifically in the Andalusian samples. Supplementary Table S2 shows the non-synonymous mutations found in the samples under study (Supplementary Table S1) with respect to the reference ON563414 with the genomic coordinates of NC_063383 [9]. The most common mutation occurs in 14 Andalusian MPXV isolates in the protein *A0A7H0DNG7*, which belongs to the Bcl-2-like protein family, which function as immunomodulators to evade the host innate immune response through the inhibition of apoptosis or blocking the activation of pro-inflammatory transcription factors [10]. Another frequent mutation shared by 13 Andalusian MPXV occurs in *A0A7H0DN47*, a transmembrane protein of unknown function. Other proteins that have been found mutated in 11 MPXV isolates are *A0A7H0DN82*, an envelope protein which has been described as a late gene transcription factor VLTF-4 [11] and *M1LBQ5*, shared by 11 Andalusian MPXV isolates, which is part of a large complex required for early virion morphogenesis [12]. Also, *A0A7H0DN66*, a component of the entry fusion complex (EFC), which consists of 11 proteins and mediates entry of the virion core into the host cytoplasm and *A0A7H0DNF5*, a soluble interferon-gamma receptor-like protein, were found mutated in 8 and 6 Andalusian MPXV isolates respectively. Thus, some differences in the immune response or in the viral replication could characterize currently circulating Andalusian isolates. There are also 96 more mutations, most of them private mutations of specific MPXV isolates and a few of them shared by up to 5 isolates as much (See Supplementary Table S2). Among them, it is worth mentioning those found in genes *A0A7H0DNG4* (in one isolate) and *A0A7H0DNG6* (in four isolates), which were previously identified as virulence genes B19R and B21R, respectively, by comparing isolates of two outbreaks in Nigeria with different mortality rates [11].

**Figure 2.**
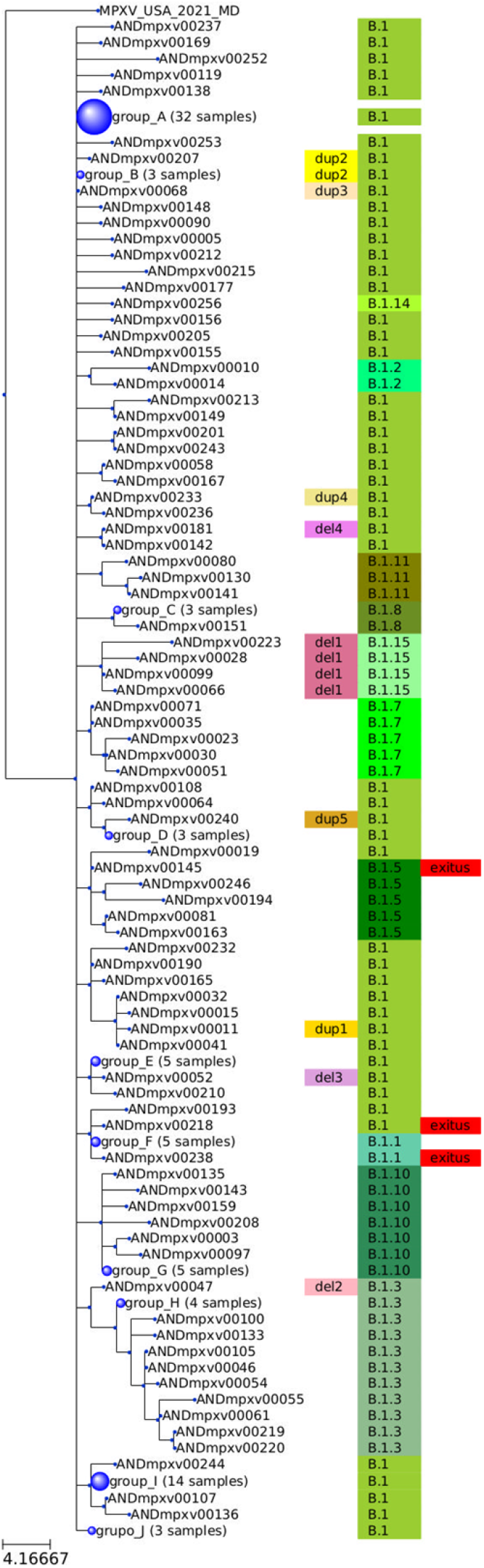
Summarized phylogeny of the Andalusian isolates, with identical sequences grouped in nodes with a size proportional to the number of corresponding isolates. Each sequence is labeled with its clade (B.1 and derived clades), and, in case of harboring a structural variation, it is also labeled. The three sequences, corresponding to the two deceased patients are labeled as well. The sequences in the nodes are: group_A: ANDmpxv00247, ANDmpxv00216, ANDmpxv00029, ANDmpxv00128, ANDmpxv00152, ANDmpxv00103, ANDmpxv00245, ANDmpxv00249, ANDmpxv00161, ANDmpxv00222, ANDmpxv00171, ANDmpxv00248, ANDmpxv00110, ANDmpxv00007, ANDmpxv00157, ANDmpxv00084, ANDmpxv00016, ANDmpxv00164, ANDmpxv00217, ANDmpxv00255, ANDmpxv00178, ANDmpxv00025, ANDmpxv00139, ANDmpxv00184, ANDmpxv00188, ANDmpxv00033, ANDmpxv00027, ANDmpxv00096, ANDmpxv00250, ANDmpxv00140, ANDmpxv00241, ANDmpxv00060; group_B: ANDmpxv00024, ANDmpxv00013, ANDmpxv00085; group_C: ANDmpxv00196, ANDmpxv00235, ANDmpxv00214; group_D: ANDmpxv00132, ANDmpxv00067, ANDmpxv00146]; group_E: ANDmpxv00075, ANDmpxv00022, ANDmpxv00048, ANDmpxv00076, ANDmpxv00123], group_F: ANDmpxv00124, ANDmpxv00154, ANDmpxv00153, ANDmpxv00021, ANDmpxv00254; group_G: ANDmpxv00251, ANDmpxv00056, ANDmpxv00095, ANDmpxv00121, ANDmpxv00020; group_H: ANDmpxv00069, ANDmpxv00162, ANDmpxv00137, ANDmpxv00017; group_I: ANDmpxv00018, ANDmpxv00074, ANDmpxv00077, ANDmpxv00126, ANDmpxv00092, ANDmpxv00242, ANDmpxv00089, ANDmpxv00098, ANDmpxv00008, ANDmpxv00172, ANDmpxv00031, ANDmpxv00147, ANDmpxv00150, ANDmpxv00012; group_J: ANDmpxv00175, ANDmpxv00102, ANDmpxv00053.

### Structural variation spectrum of the Andalusian outbreak

Among the 160 sequenced samples analyzed, a total of 15 isolates displayed structural variations (see Figure 3). Within them, 7 isolates exhibited distinct types of deletions, with sizes ranging from 912 to 6472 base pairs. Notably, the most prevalent deletion type, observed in 4 isolates, involved the partial deletion of the *A0A7H0DMZ9* protein, specifically affecting the region spanning 12143-13055 nucleotides (see del1 label in Figure 2). This protein mediates the ubiquitination and subsequent proteasomal degradation of NF-kappa-B by targeting NF-kappa-B RELA subunit to the SCF E3 ligase complex. Ubiquitination and proteasomal degradation are cellular mechanisms that are known to be targeted or modulated by some viruses to manipulate host cell signaling pathways, evade immune responses, or regulate viral protein stability [13]. Other 8 isolates bear genomic rearrangement in which one terminal part of the genome was deleted and replaced by an inverted duplication of the other end of the genome. Four of these isolates exhibit a deletion of the 3’ region and an inverted duplication of the 5’ region. This rearrangement causes the partial loss of the *A0A7H0DNG6* protein, mentioned above as related to virulence.

**Figure 3.**
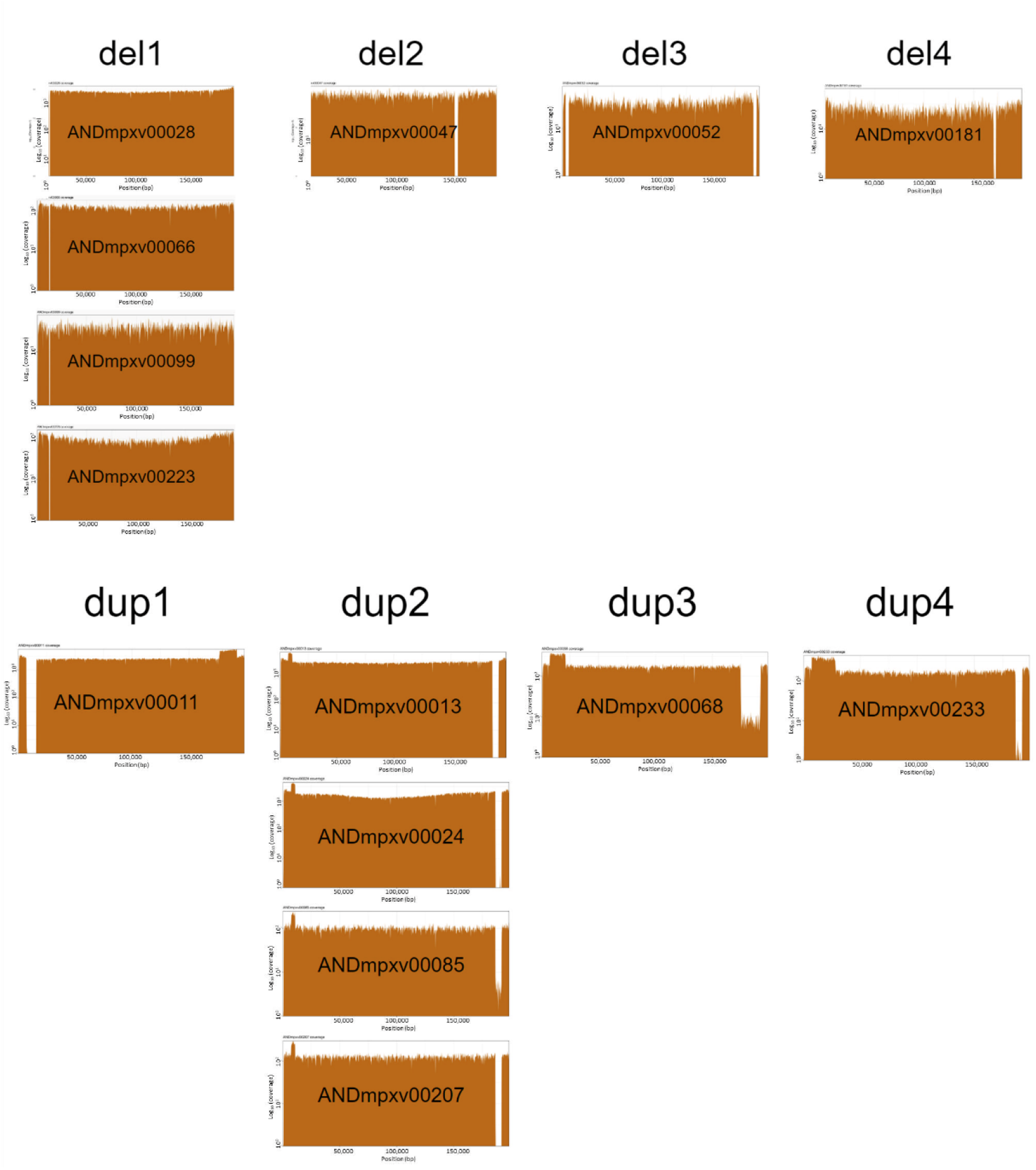
Coverage plots representing the different structural variants found.

All the structural variants seem to have appeared de novo at different points of the monkeypox phylogeny, and only in the case of del1 and dup2 (see Figure 2) seems to have configured clusters of monkeypox isolates sharing the specific variant.

### Monkeypox virus from deceased patients

Here we describe the cases of two deceased patients, represented by the sequence ANDmpxv00145, isolated from a patient with no comorbidities, and the sequences ANDmpxv00218 and ANDmpxv00238, which are two consecutive sequencing samples from an immunocompromised patient.

In the first case, the isolate ANDmpxv00145 did not present any remarkable mutation that justifies a higher virulence and, actually, it is identical to the reference sequence, except for a nucleotide synonymous mutation (C70780T) in the gene *A0A7H0DN61*, a putative nuclease. Considering both the synonymous nature of the mutation and the functional role of the implicated gene, it is improbable that this mutation alone could contribute to increased virulence in the isolate. Moreover, this synonymous mutation is also present in other two Andalusian MPXV isolates (ANDmpxv00081 and ANDmpxv00163), as well as in isolates from other countries such as Portugal, Italy, Switzerland, and Germany, all of which belong to the same clade as ANDmpxv00145. It has also been found in another Andalusian isolate (ANDmpxv00019) as a private mutation (Figure 2). To the best of our knowledge, no instances of mortality or severe complications have been documented in association with any of these isolates, thus substantiating the neutral nature of this mutation.

On the other hand, the second patient, who exhibited immunocompromised status, underwent sequencing on two occasions. During the initial sequencing, prior to the administration of tecovirimat antiviral treatment, two mutations were identified: OPG094:R194H and OPG205:E452K. The first mutation was observed in the A0A7H0DN66 gene, and has also been found in several Andalusian isolates with no apparent pathological phenotype. However, in the second sequencing conducted after tecovirimat treatment, the OPG205:E452K mutation was no longer detected, while the OPG057:A290V mutation emerged. Interestingly, this latter mutation was observed in the A0A7H0DN30 gene, whose homologous gene in Vaccinia, that present a high similarity of 99.46%, has been recognized as a target for tecovirimat [14].

On the other hand, no deletions or rearrangements of any kind of structural variation were found in any of the isolates from both deceased patients.

## Conclusions

The genomic changes detected in this study are important in assessing the microevolution of the circulating virus, although the functional impact of these mutations is still difficult to assess in the general context of virus circulation. In conclusion, the genomic surveillance platform currently running in Andalusia, created as a response of the COVID-19 pandemic, has enabled an extremely rapid response to monitor the spread and evolution of MPXV in the region, and to contribute to national and international genomic surveillance, in order to provide data and knowledge to monitor MPXV epidemics. Specific genomic surveillance criteria must be established at the national and international levels to optimize resources and increase the usefulness of its results for the control of MPXV transmission.

## Methods

### Samples

Since the MPXV appeared for the first time in the South of Spain (Andalusia), in May 26th, a few days later than the first report in Spain [15], until the end of December a total of 160 MPXV complete genomes were sequenced in the Genomic Surveillance Circuit of Andalusia [6, 7]. Supplementary Table S1 lists the viral genomes sequenced. The isolates were evenly sampled across all Andalusia.

### Sequencing method

Prior to DNA extraction from ulcerative lesion samples using the QIAamp DNA kit (Qiagen), sonication and DNAse/RNAse treatment was performed [8]. Subsequently, shotgun metagenomics was performed. In brief, DNA libraries were prepared using the Illumina DNA Prep kit (Illumina, San Diego, CA, USA) and IDT for Illumina DNA/RNA UD Indexes sets (Ilumina). The quality of the libraries was validated by Qubit 4 fluorometer (Thermo Fisher Scientific, Waltham, MA, USA). Sequencing was performed on Nextseq 550/1000 (Illumina).

### Data processing

Sequencing data were analyzed using in-house scripts and the nf-core/viralrecon pipeline software [16], version 2.4.1. Briefly, after read quality filtering, sequences for each sample are aligned to the high quality MPXV isolate OX044336.2 [17] related to the 2022 outbreak using bowtie2 algorithm [18]. Genomic variants were identified through iVar software [19], using a minimum allele frequency threshold of 0.25 for calling variants and a filtering step to keep variants with a minimum allele frequency threshold of 0.75. Using the set of high confidence variants and the OX044336.2 genome, a consensus genome per sample was finally built using bcftools [20].

### Phylogenetic analysis

Phylogenetic analysis was carried out on the obtained MPXV genomes in the context of a world-wide representative set of MPXV genomes available in NCBI virus [21] and virological.org using the Augur application [22], whose functionality relies on the IQ-Tree software [23]. The MAFFT program [24], was utilized for the multiple alignment, using the isolate MPXV-M5312_HM12_Rivers (NC_063383.1) as reference. A maximum likelihood method with a general time reversible model with unequal rates and unequal base frequencies [25] was used to reconstruct the viral phylogeny. The results can be viewed in the Nextstrain Auspice [26] local server, which is now part of the Genomic Surveillance Circuit of Andalusia [27].

The amino acid substitutions for each of the 160 samples sequenced in Andalusia (Supplementary Table S1) were obtained using the nextclade web application [28]. Specifically using as pathogen reference “Human Monkeypox Clade B.1” (ON563414), which belongs to the same clade as the Andalusian samples but in the coordinates of the isolate NC_063383.

### Structural variations

To identify samples that could harbor large structural variants or deletions, analyses of coverage plots were conducted. Specifically, samples exhibiting regions of low coverage or regions with high coverage in the coverage plots were located [29].

To confirm potential structural variants, reads from the samples displaying non-homogeneous patterns in the coverage plots were aligned using the bwa program [30], employing the “-a” argument to retain all alignments. The sample OX044336.2 was used as the reference in this process.

The determination of the breakpoints for deletions and the insertion points for rearrangements was achieved by selecting reads that did not have an appropriate insert size. This indicated that the mate pairs mapped to both sides of a deletion or different regions in the case of duplication/rearrangement. To filter and obtain these reads, the samtools application [31] was used with the “-F14” filter, which eliminates reads that have not mapped and properly mapped reads. Furthermore, to confirm the start and end points, only reads with chimeric alignments, where a portion of the read aligned to one genomic region and another portion aligned to a different region, were retained. This selection was made using the flag “2048” to filter the reads.

Finally, in IGV [32], it was confirmed that the accumulation of reads with inappropriate insert sizes and chimeric reads corresponded to the positions where the coverage plot exhibited changes.

## Data availability

The 160 MPXV genomes are available at the European Nucleotide Archive (ENA) repository under the accession number PRJEB55075. Supplementary Table S1 provides also each individual sample ENA IDs.

## Acknowledgements

This work was supported by Spanish Ministry of Science and Innovation (grants PID2020-117979RB-I00 and FJC2021-046546-I), the Instituto de Salud Carlos III (ISCIII), co-funded with European Regional Development Funds (ERDF) (grant IMP/00019), it has also been funded by Consejería de Salud y Consumo, Junta de Andalucía (grants COVID-0012-2020), and by grant ELIXIR-CONVERGE - Connect and align ELIXIR Nodes to deliver sustainable FAIR lifescience data management services (AMD-871075-16), funded by EU – H2020.

## Supporting information

**Supplementary Table S1**. Monkeypox samples sequenced in this study, along with their collection date, origin and individual ENA IDs, belonging to the collective accession number PRJEB55075.

**Supplementary Table S2**. Non-synonymous mutations found in the samples under study with respect to the reference ON563414 in NC_063383 coordinates.

**Supplementary Figure 1**. Whole phylogeny of the Andalusian isolates, with identical sequences grouped in nodes with a size proportional to the number of corresponding isolates. Each sequence is labeled with its clade (B.1 and derived clades), and, in case of harboring a structural variation, it is also labeled. The three sequences, corresponding to the two deceased patients are labeled as well.

